# Multi-stage chemical heating for instrument-free biosensing

**DOI:** 10.1101/367029

**Authors:** John P. Goertz, Kenya M. Colvin, Andrew B. Lippe, John L. Daristotle, Peter Kofinas, Ian M. White

**Affiliations:** Fischell Department of Bioengineering and University of Maryland, College Park, USA; Department of Chemical and Biomolecular Engineering, University of Maryland, College Park, USA

## Abstract

Improving the portability of diagnostic medicine is crucial to alleviating global access-to-care deficiencies. This requires not only designing devices that are small and lightweight but also autonomous and independent of electricity. Here, we present a strategy for conducting automated multi-step diagnostic assays using chemically generated, passively regulated heat. Ligation and polymerization reagents for Rolling Circle Amplification of nucleic acids are separated by melt-able phase-change partitions, thus replacing precise manual reagent additions with automated partition melting. To actuate these barriers and individually initiate the various steps of the reaction, field ration heaters exothermically generate heat in a thermos while fatty acids embedded in a carbonaceous matrix passively buffer the temperature around their melting points. Achieving multi-stage temperature profiles extends the capability of instrument-free diagnostic devices and improves the portability of reaction automation systems built around phase-change partitions.

---

Access to healthcare remains one of the primary challenges of modern medicine. Technological advances often remain concentrated in wealthy urban centers, out of reach to rural and poor populations in developing and developed nations alike. Alleviating this disparity is not simply a matter of making existing techniques affordable: many traditional assay platforms are incompatible with field-use at any cost. Instead, alternative technologies must be designed for high portability (small, lightweight, not reliant on electricity) and ease-of-use (simple and autonomous, with minimal hands-on steps).

Two primary approaches have emerged to answer this need: chip- and paper-based microfluidics.^1,2^ Traditional microfluidic devices that utilize micro-fabricated fluidic networks are capable of housing numerous reactions with myriad components that proceed in a well-orchestrated pattern, yet the requisite pumps and other peripheral equipment severely impair portability.^3^ While paper devices significantly improve the portability of biosensing reactions, the simplicity that makes them easy to use limits the throughput and complexity of assays they can support.^4^ To address this gap between miniaturized assays suitable only for laboratory use and those restricted to field use, we recently described the novel approach of employing thermally-reversible barriers to sequester reagents within a common PCR tube.^5^ This approach demonstrated the potential to offer the tight reaction control of microfluidics with portability and ease-of-use that parallels paper devices.

These “phase-change partitions” consist of ordinarily-solid purified hydrocarbon waxes that exhibit sharply-defined melting transitions at distinct temperatures. Reagents for each step of a multi-part reaction remain isolated from one another until the respective barrier is melted, at which point a sample solution sinks through the now-molten alkane and mixes with the reagent beneath. This approach allows arbitrarily long reaction stages and at least five distinct reagent zones within a single 200 PCR tube. While the temperature range spanned by the alkanes we employed is narrow enough to remain accessible to simple heating devices, the melting transitions are discrete enough to avoid the need for tightly-calibrated temperature control. Indeed, we demonstrated actuation of these phase-change partitions in a simple water-bath as well as a commercial thermocycler.

However, even a temperature-regulated water-bath requires a consistent source of electricity, unavailable in field settings or low-resource clinics. A similar challenge is faced by many isothermal nucleic acid amplification techniques such as LAMP (loop-mediated isothermal amplification)^6,7^ and RPA (recombinase-polymerase amplification)^8,9^, which require elevated temperatures to achieve highly-sensitive detection of pathogens. Numerous groups have employed chemically-generated heat with thermal buffers to reach the incubation temperature for these reactions.^10–12^ This is typically achieved using the exothermic hydration of calcium oxide or the galvanic corrosion of MgFe alloys in the presence of saline.^12,13^ This latter reaction is extensively employed in military Meal-Ready-to-Eat (MRE) field ration heaters and thus has been thoroughly optimized to rapidly reach boiling temperatures while remaining compact and lightweight.

To achieve prolonged incubation at temperatures amenable to biochemical reactions rather than a brief burst of excessive heat, researchers have employed phase-change materials (PCMs) as thermal buffers in these exothermic systems.^10–16^ These materials surround the reaction compartment so that, as they melt, the temperature of the reaction remains near that of the compound’s melting point (Figure S-1, Supplementary Material).^17^ Phase-change materials with specified thermal characteristics are commercially available^18^ but can also be inexpensively fashioned from materials such as fatty acids and hydrated salts with high latent heats of fusion and desirable melting temperatures.^19,20^ Previous reports have described systems which are designed for only a single operating temperature; here, we present the use of MRE heaters with blended PCMs to achieve multi-stage temperature profiles. We leveraged this platform to sequentially actuate two phase-change partitions in a PCR tube, simultaneously providing ideal operating temperatures for the ligation and polymerization stages of Rolling Circle Amplification (RCA). Our results demonstrate the potential for platforms based on phase-change partitions to automate the field use of complex, multi-stage biosensing reactions without the need for electricity.

## Materials and Methods

### Materials

Carbon black (CB) (99.9%), lauric (dodecanoic) acid (LA) (98%), and palmitic (hexadecanoic) acid (PA) (95%) were purchased from Alfa Aesar (Haverhill, MA). Sodium thiosulfate pentahydrate, sodium acetate trihydrate, and carboxymethyl cellulose (CMC) were purchase from Sigma. 0.2 mL high-profile PCR tubes were purchased from USA Scientific (Ocala, FL). RCA reagents were purchased from New England Biolabs (Ipswitch, MA), and DNA sequences were purchased from Integrated DNA Technologies (Coralville, IA). DNA Sequences used are:

Trigger: 5’-TAG TCG AGA CAT CCG AGA CA-3’

Template: 5’-Phos-GTC TCG ACT AAA AAC CCA ACC CGC CCT ACC CAA AAG AGA CAT CCG TTT TGT CTC GGA T-3’

MRE Heaters were provided by Luxfer Magtech Inc. Tahoe Trails 10 oz vacuum insulated double wall stainless steel travel tumblers, 4 × 5 inch sealable tea bags, Kayose natural tea filter bags, and the iTouchless handheld heat bag sealer were purchased from Amazon. 1 mm nylon mesh sieves were purchased from Component Supply Company. The custom-designed thermos insert was 3D printed in ABS with a Zortrax M200; the .stl file can be found in the Supplementary Material. All remaining materials were purchased from MilliporeSigma (Burlington, MA).

### Preparation of Encapsulated PCMs

Fatty acids and hydrated salts were melted on a hot plate under magnetic stirring. Carbon black or activated carbon was mixed with melted fatty acids at specified weight ratios. The hydrated salts were mixed first with 5 wt% CB and subsequently with 10 wt% CMC; a small amount of methanol was added to allow CB to mix with the molten salt hydrate. The resulting pastes were spread on aluminum foil to cool, ground in a mortar and pestle, and sieved to obtain granules <1 mm in diameter. To produce systems with multiple temperature stages, multiple PCMs were encapsulated separately then mixed after cooling so as not to impact their individual melting points.

### Form-Stability of Encapsulated PCMs

To assess the stability of the encapsulated PCMs against melt leakage, ~4 g composite was placed in Kayose tea bags, sandwiched between paper towels and then aluminum foil, and placed on an 80 °C hotplate. Mass of the composite was taken before and after one hour of incubation.

### Differential Scanning Calorimetry (DSC)

Approximately 10 mg samples were sealed in aluminum hermetic pans (TA Instruments) using a sample encapsulation press. DSC measurements were made on a TA Instruments DSC Q100. Samples were held isothermal at 0 °C for 5 min, then heated to 100 °C and cooled to 0 °C at a rate of 3 °C min^−1^, ± 0.20 °C amplitude, with a modulation period of 60 s for two continuous cycles.

**Figure 1.**
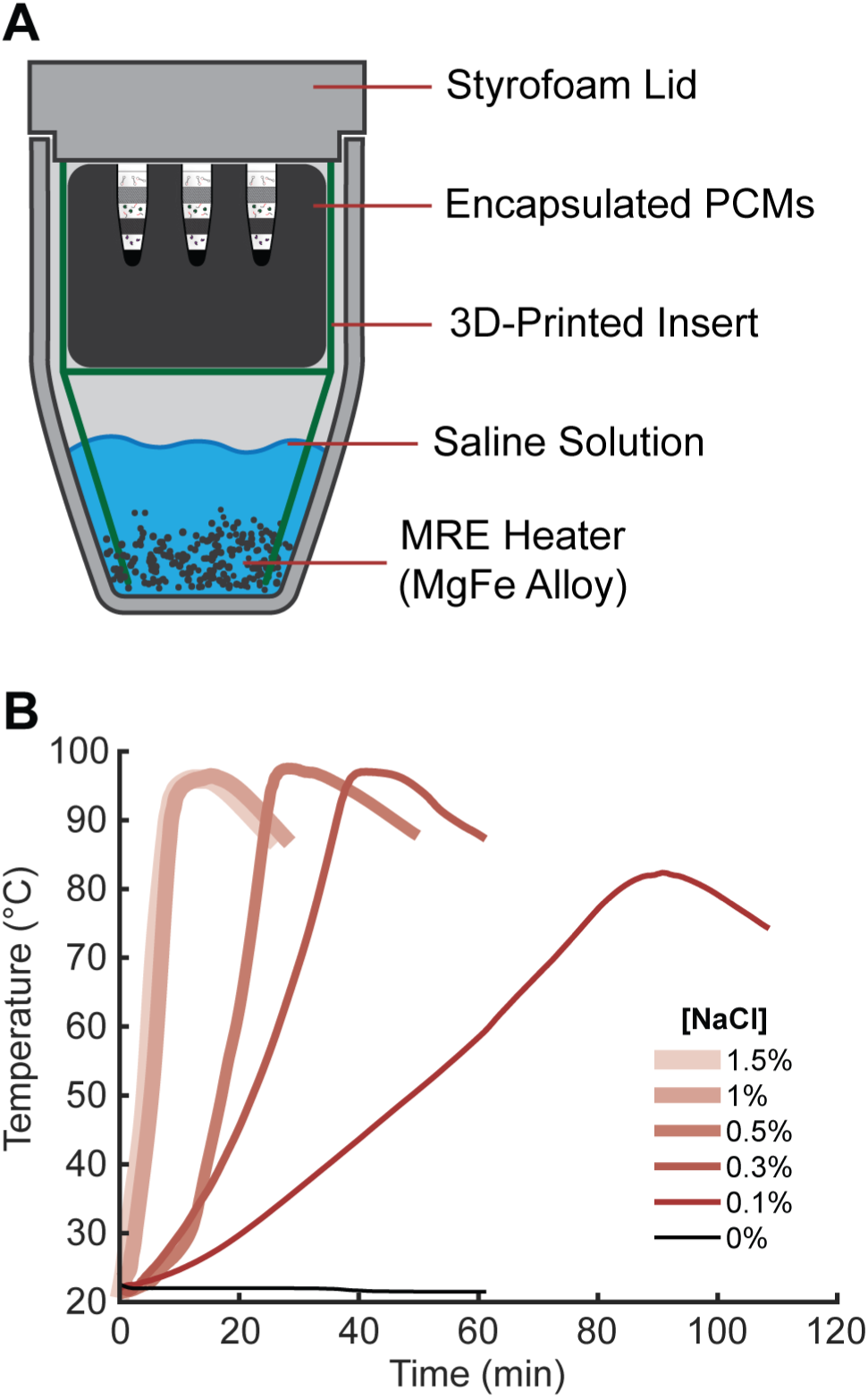
Chemical heating. **A)** A 10 oz thermos provides the housing for PCM-MRE actuation of phase-change partitioned assays. **B)** Decreasing the saline concentration used to initiate the exothermic reaction provides a more gradual temperature profile, facilitating passive thermal regulation with PCMs.

### Chemical Heating

MRE heaters were used as is to determine the temperature profile of the exothermic reaction between saline and the MgFe alloy in the MREs. A single packet of the MgFe alloy was added to a thermos followed by 100 mL of saline. The reaction was examined using NaCl concentrations of 0, 0.1, 0.3, 0.5, 1.0, and 1.5 wt%. Temperature was recorded using a Sparkfun waterproof temperature sensor (DS18B20) and an Arduino Uno.

### Passively Regulated Temperature

MRE heaters were repackaged using heat-sealable tea bags, as described previously. MgFe alloy granules from MREs were distributed into each tea bag in 3.70 g allotments, sealed, then placed in the bottom of the vacuum thermos. The exothermic reaction was initiated by adding 100 mL of 0.1% saline, after which a 3D-printed insert made of ABS was placed in the container and covered with aluminum foil, suspending the PCM above the saline level. A Styrofoam lid provided insulation at the top of the thermos, while a small hole allowed hydrogen gas produced in the chemical reaction to vent. The temperature within the PCM was recorded using a Vernier Labquest Mini and a stainless steel temperature probe.

### Phase-Change Partitioned Assays

The pH indicator demonstration was constructed in a PCR tube, from top to bottom, with 40 μL indicator solution (1 mM H_2_SO_4_, 1% Triton, 60% Indicator), 50 μL octadecane, 10 μL each buffer A (800 mM NH_4_OAc, 250 mM Tris, pH 6.0) and B (5 M NH_4_OAc, 500 mM Tris, pH 8.0), 40 μL tetracosane, and 30 μL 9.8 M NH_4_OH. The RCA reaction was constructed, from top to bottom, with 20 μL 1.36 μM Template DNA in DI water, 50 μL octadecane, 20 ligase solution (18 U/μL T4 DNA Ligase, 1.8× Ligase Buffer with ATP), 50 tetracosane, 20 μL polymerase solution (1.1 U/μL Bst 3.0 DNA Polymerase, 2.7× Isothermal Amplification Buffer, 2.7 mM dNTPs, 1.36 μM Trigger DNA). Tubes were embedded into a mixture of 20 g encapsulated LA and 20 encapsulated PA, then heated as described above with an MRE heater initiated by 0.1% saline. Reactions marked *L* in Figure 4 were removed once the vessel temperature reached 40 °C, and reactions marked *F* were removed one hour after the vessel temperature exceeded 55 °C.

### Gel Electrophoresis

Denaturing polyacrylamide was used to analyze RCA products. Reactions were removed at the respective time and halted by immersion in ice water. Reaction solutions were extracted, mixed with two parts 12 M urea, heated to 95 °C for five minutes, then run on a 15% gel at 500V for 15 minutes in a BioRad mini PROTEAN.

## Results

The usability of phase-change partitions for diagnostic reactions in low resource settings heavily depends on being able to easily manipulate the heat source without the use of electricity or additional equipment. Our multi-stage heating device consisted of an off-the-shelf vacuum thermos separated by a 3D-printed insert: a lower chamber contained the MRE alloy packet while the encapsulated PCMs and reaction tubes were housed in an upper chamber above the saline level (Figure 1A). We were able to easily fine tune the temperature profile of the saline-activated MgFe alloy by changing the concentration of salt in the solution (Figure 1B). The saline pack included with the MREs contained 1.5 wt% salt and caused a rapid increase in temperature up to 97°C. Reducing the salt concentration increased the time it took for the MgFe alloy to reach its maximum temperature while lowering that maximum, reducing the thermal burden needed to be buffered by PCMs.

We investigated two fatty acids (LA, m.p. ~43 °C, and PA, m.p. ~63 °C) and two hydrated salts (sodium thiosulfate pentahydrate, Na_2_S_2_O_3_·5H_2_O, m.p. ~48 °C; and sodium acetate trihydrate, NaOAc·3H_2_O, m.p. ~58 °C) for use as PCMs. Ideally, PCMs must be encapsulated to prevent leakage of the melted material during operation and, in the case of hydrated salts, to prevent phase-separation in the molten state; doing so also has the advantage of improving the thermal conductivity of the material. There is an extensive body of literature devoted to such encapsulation techniques for the purposes of solar heating as well as “smart” construction and textile materials, most of which entail either formation of core-shell microparticles or distribution of the PCM within a porous matrix.^21^ Here, we chose carbon black (CB) as an encapsulant for fatty acids due to its affordability, ease of encapsulation, and thermal-conduction properties. The fatty acid was melted, mixed rapidly with CB to penetrate the porous matrix, cooled, ground, and sieved (Figure 2A). Upon subsequent re-melting, surface tension caused the molten fatty acid to remain entrapped within CB pores. This composite exhibited bulk minimal leakage at elevated temperatures when the CB mass fraction was 20% or greater (Figure 2B); curiously, CB provided greater form-stability than activated carbon, despite the latter’s nominally higher surface area to volume ratio and prominent position in the PCM literature.^19,22^ For hydrated salts, 5% CB with 10% CMC achieved adequate form-stability.^22^

**Figure 2.**
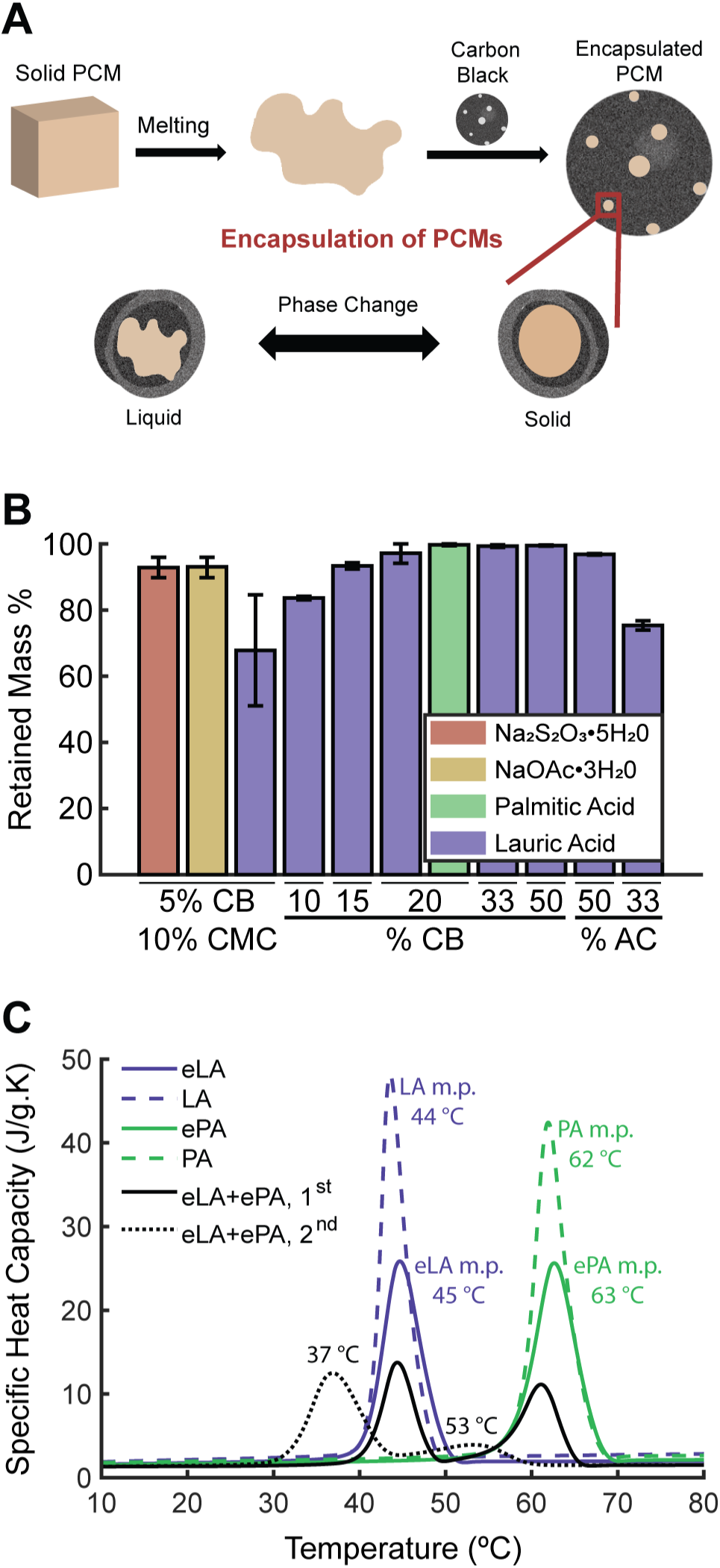
PCM Encapsulation. **A)** Fatty acids are melted then mixed with carbon black, causing the molten PCM to impregnate the pores of the carbon matrix. Upon re-melting, capillary tension prevents the PCM from leaking out of the encapsulation. **B)** Form-stability of encapsulations was evaluated by measuring the mass lost after one hour of heating at 80 °C. **C)** DSC analysis demonstrates the narrow melting profile of fatty acids. LA, PA: pure fatty acid alone. eLA, ePA: fatty acid with 20% CB. eLA+ePA: 1:1 mixture of each fatty acid individually encapsulated in 20% CB, first and second run. Black text labels refer to eLA+ePA, 2^nd^ peaks.

**Figure 3.**
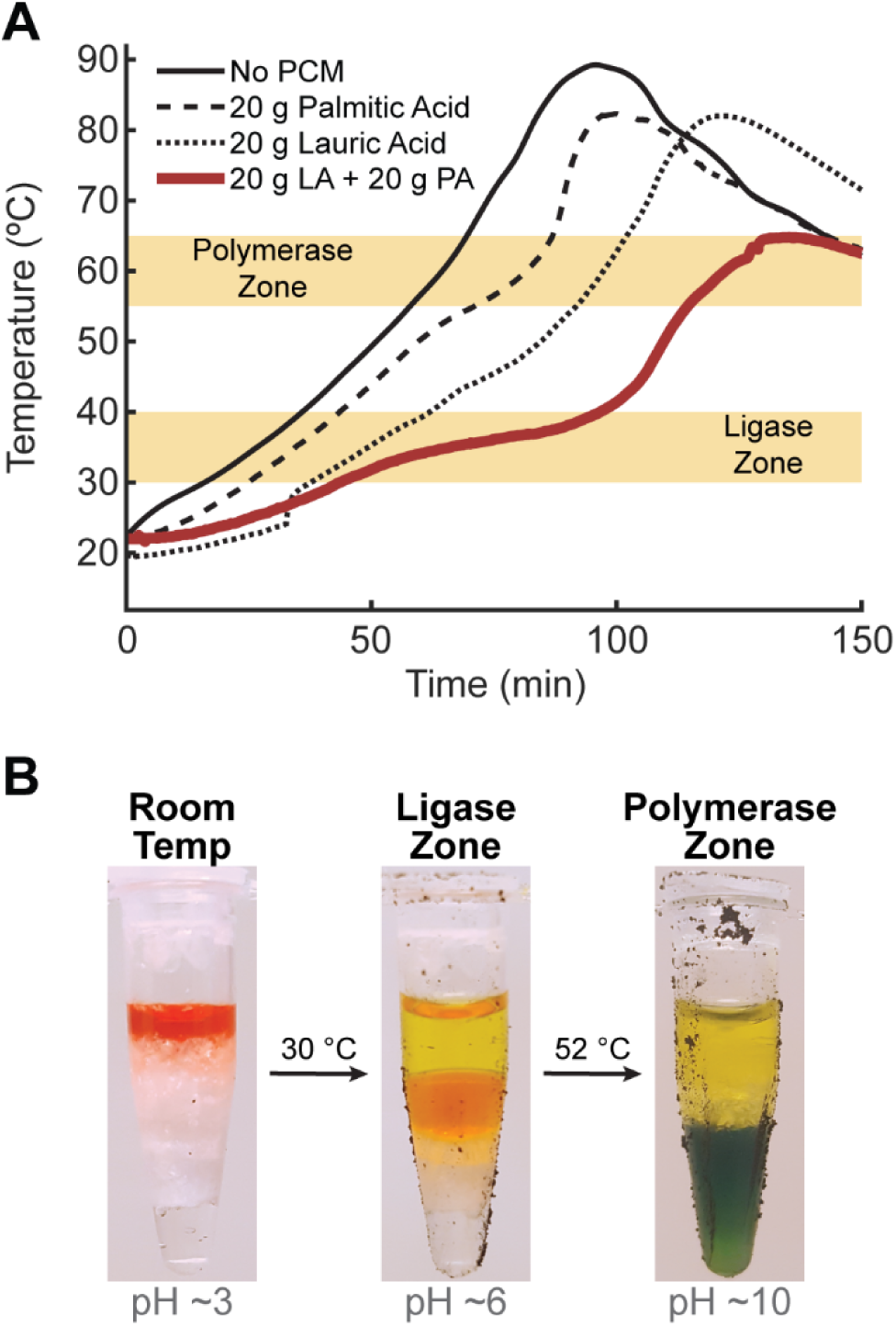
Multi-stage chemical heating. **A)** An MRE heater activated with 0.1% saline and buffered with a mixture of 20 g encapsulated lauric acid and 20 g encapsulated palmitic acid produces two distinct temperature zones amenable to different biochemical processes. **B)** This two-stage heating can be used to actuate phase-change partitions. Here, pH Indicator is stepped through different buffers sequestered by eicosane and tetracosane barriers.

We used differential scanning calorimetry (DSC) to investigate the thermal properties of our encapsulated PCMs. As shown in Figure 2C and Figure S-2 (Supplementary Material), encapsulation resulted in minimal change in PCM melting point (defined as the temperature at maximal specific heat capacity), implying no chemical interaction between core and matrix materials. We also examined a mixture of encapsulated LA and encapsulated PA, which displayed significantly lower melting points when melted a second time. This suggests that the molten fatty acids migrate between the carbon black particles, mixing with one another and mutually depressing their respective melting points.

This mixture of encapsulated fatty acids provided an adequate thermal buffer for to produce multi-stage temperature profile from an MRE heater. By combining 20 g encapsulated PA with 20 g encapsulated LA, the temperature profile generated by an MRE heater with 0.1% saline was successfully modulated to exhibit an approximately 1 hour hold between 30 and 40 °C and a greater than 1 hour hold between 55 and 65 °C (Figure 3A). This temperature profile allowed well-controlled actuation of phase-change partitions, demonstrated by stepping a pH indicator solution through sequential eicosane and tetracosane barriers to mix with various buffers (Figure 3B). Combinations of encapsulated hydrated salt also produced multiple temperature stages (Figure S-3, Supplementary Material).

The lowered melting points of the encapsulated fatty acid mixture provided ideal temperature regimes to achieve Rolling Circle Amplification. RCA is a two-step method for DNA detection: a template sequence is first ligated into a circle, then a complementary trigger sequence is extended by a polymerase to continuously replicate the template.^23,24^ While each stage requires 30-60 minutes at only a single temperature, the ligase enzyme is most active between 30 and 40 °C and the fastest polymerase enzymes are active between 55 and 65 °C; furthermore, the two steps must be performed separately, since premature extension of the trigger sequence along an un-circularized template prevents ligation and continuous amplification.

**Figure 4.**
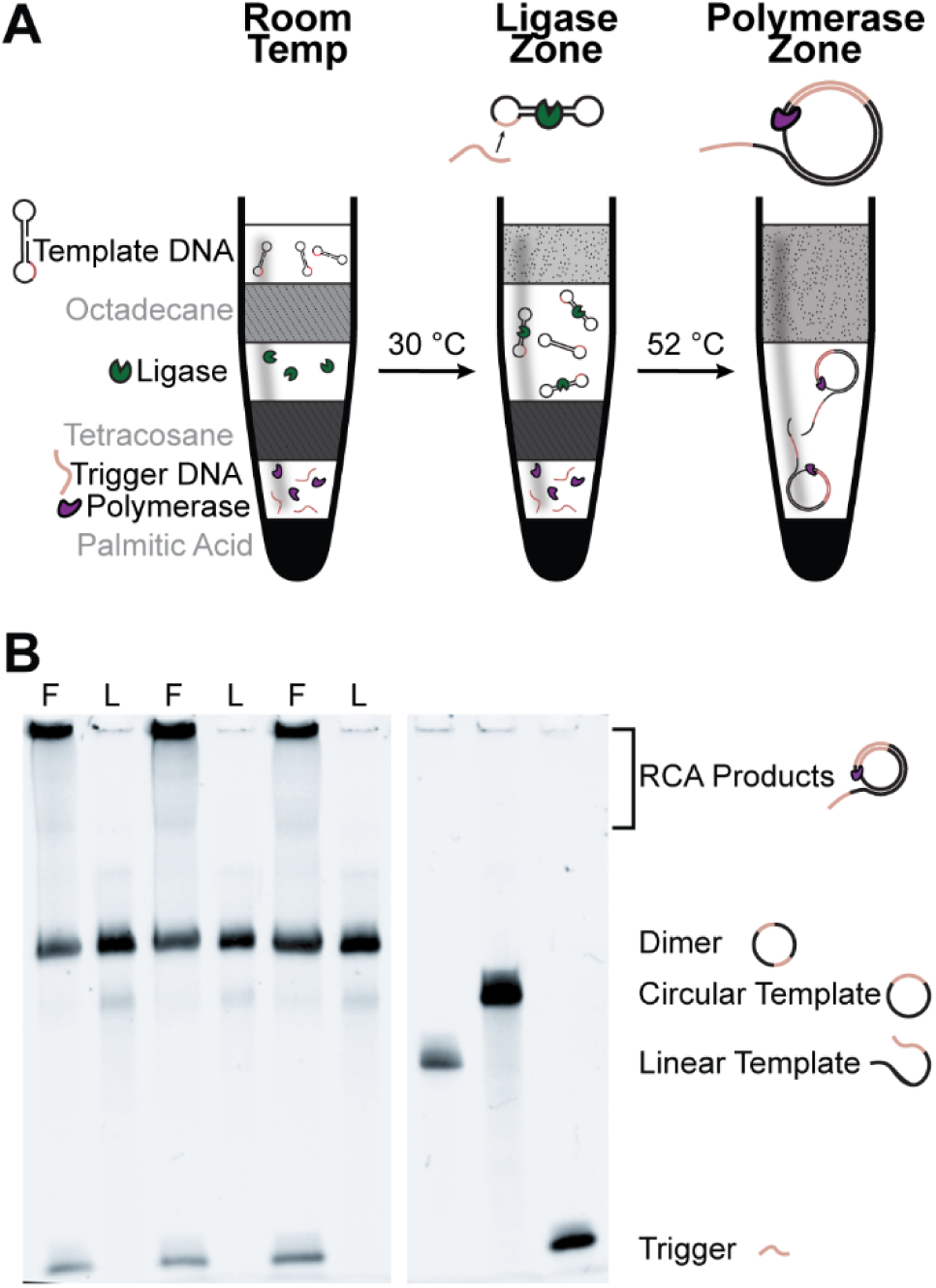
Portable biosensing with multi-stage chemical heating. **A)** The phase-change partitioned RCA assay is initiated by melting of an octadecane layer once the tube exceeds 30 °C, causing ligase enzyme to ligate the template DNA into a circle. The amplification stage is initiated by melting of a tetracosane layer once the tube exceeds 52 °C, at which point the polymerase extends a trigger sequence to continuously replicate the template. **B)** Denaturing acrylamide gel electrophoresis reveals successful ligation in all reactions, and successful generation of amplicon in those incubated for the full duration (*F*). The absence of trigger DNA in reactions incubated only for the ligase portion (*L*) confirms the integrity of the tetracosane barrier below its melting point. Note that the apparent difference in circular template band intensity between Ligase-only and Full reactions is due to the further dilution by the polymerase solution in the latter.

We constructed a partitioned RCA reaction by placing a dumbbell-forming template DNA sequence above a layer of octadecane (m.p. 30 °C), followed by a buffer containing ligase, a layer of tetracosane (m.p. 52 °C), and finally a buffer containing trigger DNA and polymerase (Figure 4A). Six such reactions were run in parallel in the MRE-PCM system described above. Three were removed once the thermos reached 40 °C and the remaining three were incubated further until the thermos had spent an hour above 55 °C, during which time the temperature never exceeded 65 °C. As demonstrated by gel electrophoresis (Figure 4B), both ligation and polymerization proceeded efficiently; furthermore, the trigger sequence is not present in the reactions incubated only until 40 °C, indicating that the phase-change partitions completely sequestered the various reaction components.

## Conclusion

We have demonstrated the electricity-free automation of a multi-step biosensing reaction. The phase-change partition platform reported previously enabled stable separation of reactants with thermally-reversible alkane barriers; the current work provides a system capable of actuating these partitions in an automated, field-compatible manner. Tempering the saline concentration added to MRE heater granules metered the rate of the accompanying exothermic reaction, while phase-change materials buffered the reaction temperature within multiple sequential ranges amenable to biochemical reactions. Encapsulating the PCMs within porous matrices prevented bulk leakage, enabling re-use. When used alone, encapsulated LA produced a temperature hold of approximately 45 °C, but when combined with encapsulated PA, the first temperature hold occurred at a temperature regime more amenable to T4 DNA Ligase, between 30 and 40 °C. The melting point of PA was similarly depressed, remaining within an optimal region for *Bst* polymerase. Additionally, future investigations should explore numerical approaches to quantitatively model PCM-MRE temperature profiles and accelerate the development cycle of non-instrumented diagnostic assays.

We successfully actuated a phase-change partition system with passively-buffered chemical heating to demonstrate the capacity of this system to automate nucleic acid amplification. This report extends the compatibility of the phase-change partition platform to include not only well-equipped laboratories (via thermocyclers) and generic clinics (via water baths), but also resource-poor settings and field operation (via multi-stage PCM-MRE heaters). The key advantage of such broad compatibility is that it enables a common form factor to be employed in diverse settings: the same assay can be given to a central lab technician and a field nurse. Our results demonstrate that phase-change partitions have the potential to bridge the current gap between centralized and remote diagnostic platforms. Now further developments are necessary to adapt a wide range of clinical assays to this system and support efforts to close the urban-rural divide that persists in 21^st^ century medicine.

## Acknowledgements

We would like to thank Luxfer Magtech Inc for their kind donation of MRE heaters. We would also like to thank Alessandra Zimmermann for her comments on the final manuscript.

## Supplementary Material

### Supporting Methods

#### Fatty Acid Characterization

Each well of a PCR strip was filled with 200 μl of a pure fatty acid. A thermocouple was placed in each well and the fatty acid solidified around the thermocouple. The melting rate profile of the fatty acids were then monitored as the temperature increased to 80 °C at a constant rate of 0.1 °C/min in the thermocycler. Data were fit with a smoothing spline.

**Figure S-1.**
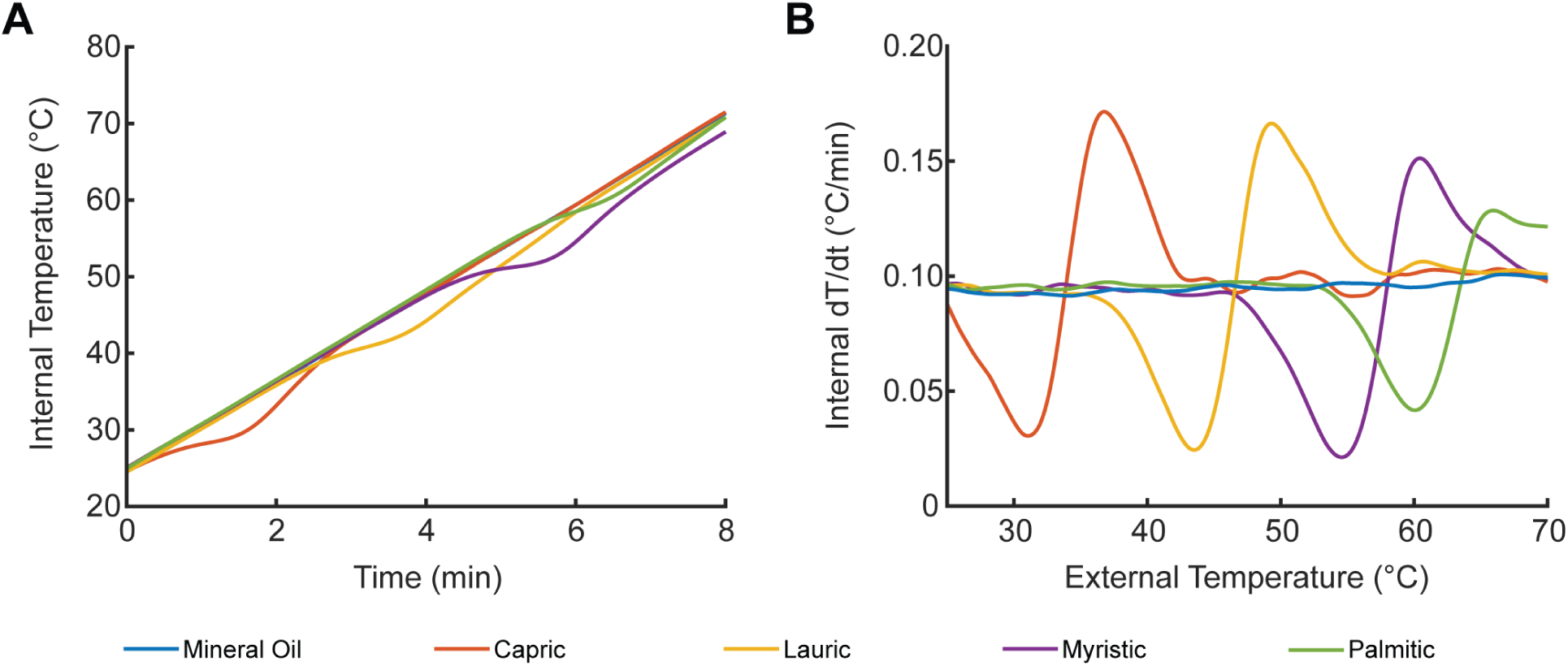
Passive temperature regulation with fatty acids. **A)** Internal temperatures and **B)** respective internal temperature vs. time derivatives of fatty acids exposed to a 0.1 °C/min temperature ramp. The fatty acids buffer the temperature around their melting point, slowing the temperature change in their interiors as they melt.

**Figure S-2.**
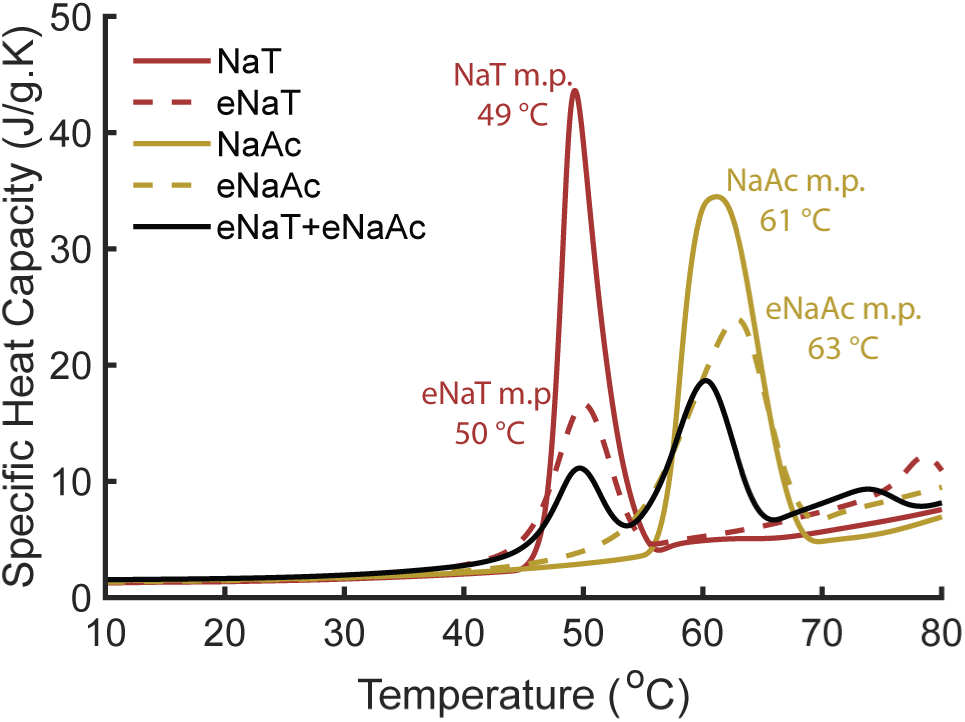
DSC analysis of hydrated salts. *NaT:* Na_2_S_2_O_3_·5H_2_O. *eNaT:* NaT with 10% CB and 5% CMC. *NaAc:* NaOAc·3H_2_O. *eNaAc:* NaAc with 10% CB and 5% CMC. *eNaT+eNaAc:* 1:1 mixture of eNaT and eNaAc.

**Figure S-3.**
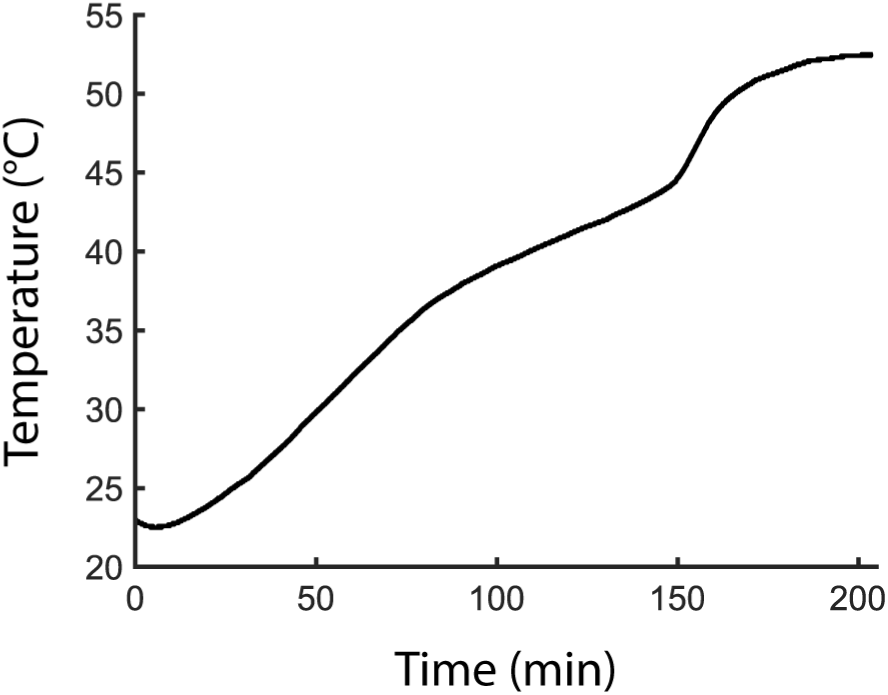
Temperature profile of 30 g Na_2_S_2_O_3_·5H_2_O (m.p. ~48 °C) and 20g NaOAc·3H_2_O (m.p. ~58 °C). Two extended temperature holds are observed, initially between 40 °C and 45 °C and subsequently between 50 °C and 55 °C.

